# Draft genome of the lowland anoa (*Bubalus depressicornis*) and comparison with buffalo genome assemblies (Bovidae, Bubalina)

**DOI:** 10.1101/2022.06.14.496191

**Authors:** Stefano Porrelli, Michèle Gerbault-Seureau, Roberto Rozzi, Rayan Chikhi, Manon Curaudeau, Anne Ropiquet, Alexandre Hassanin

## Abstract

Genomic data for wild species of the genus *Bubalus* (Asian buffaloes) are still lacking while several whole genomes are currently available for domestic water buffaloes. To address this, we sequenced the genome of a wild endangered dwarf buffalo, the lowland anoa (*Bubalus depressicornis*), produced a draft genome assembly, and made comparison to published buffalo genomes.

The lowland anoa genome assembly was 2.56 Gbp long and contained 103,135 contigs, the longest contig being 337.39 kbp long. N50 and L50 values were 38.73 kbp and 19.83 kbp, respectively, mean coverage was 44x and GC content was 41.74%. Two strategies were adopted to evaluate genome completeness: (i) determination of genomic features with *de novo* and homology-based predictions using annotations of chromosome-level genome assembly of the river buffalo, and (ii) employment of benchmarking against universal single-copy orthologs (BUSCO). Homology-based predictions identified 94.51% complete and 3.65% partial genomic features. *De novo* gene predictions identified 32,393 genes, representing 97.14% of the reference’s annotated genes, whilst BUSCO search against the mammalian orthologues database identified 71.1% complete, 11.7% fragmented and 17.2% missing orthologues, indicating a good level of completeness for downstream analyses. Repeat analyses indicated that the lowland anoa genome contains 42.12% of repetitive regions. The genome assembly of the lowland anoa is expected to contribute to comparative genome analyses among bovid species.

## 1. Introduction

The lowland anoa, *Bubalus depressicornis* (C. H. Smith, 1827), is a wild dwarf buffalo endemic to Sulawesi and Buton Islands, where it can be found in sympatry with the mountain anoa, *Bubalus quarlesi* (Ouwens, 1910). Both anoa species are currently classified as endangered with declining populations due to hunting and habitat loss (Burton et al. 2016). Because of their singular appearance, they were initially described in their own genus *Anoa* (Ouwens 1910). However, *Anoa* was not regarded as a valid genus in more recent classifications, in which both anoa species were ascribed to the genus *Bubalus*, together with the wild water buffalo–sister-group relationship between *Bubalus arnee* (Kerr, 1792) and the tamaraw - *Bubalus mindorensis* Heude, 1888 (Groves 1969; IUCN 2022). Molecular studies based on mitochondrial sequences have supported a sister-group relationship between *Bubalus depressicornis* and *Bubalus quarlesi* (Schreiber et al., 1999; Priyono et al., 2020). In addition, the mitogenome of the lowland anoa was found to be equally distant from those of the two types of domestic water buffalo, the river buffalo from the Indian subcontinent and Mediterranean countries and the swamp buffalo from China and Southeast Asia (Hassanin et al., 2012). Since the same phylogenetic pattern was recovered from the analyses of two nuclear datasets, one based on 30 autosomal genes and the other based on two genes of the Y chromosome, Curaudeau et al. (2021) have concluded the existence of two species of domestic buffaloes: *Bubalus bubalis* (Linnaeus, 1758) for the river buffalo and *Bubalus kerabau* Fitzinger, 1860 for the swamp buffalo, which diverged during the Pleistocene at around 0.84 Mya. As discussed in Curaudeau et al. (2021), the two domestic species can easily be distinguished based on coat and horn characteristics (Castelló 2016), and they have different karyotypes: *Bubalis bubalis* has 2n = 50 chromosomes with a fundamental number (FN) equal to 58; whereas *Bubalus kerabau* has 2n = 48 chromosomes and FN = 56 (Nguyen et al., 2008).

With rapid progress and cost reduction in sequencing technologies, many whole genomes of domestic bovid species have been sequenced. Whole-genome sequencing has allowed the identification of variants involved in domestication and genetic improvement for several livestock species such as cattle and buffaloes (Zimin et al., 2009; Canavez et al., 2012; Li et al., 2020; Rosen et al., 2020). Chromosome-level genome assemblies include those of the domestic cow, *Bos taurus* (Zimin et al., 2009), the domestic river buffalo, *Bubalus bubalis* (Deng et al., 2016), the swamp buffalo, *Bubalus kerabau* (reported as *Bubalus carabanensis* in Luo et al. (2020) but see Curaudeau et al. (2021) for further taxonomic information), the domestic Yak, *Bos grunniens* (Zhang et al., 2021) and the zebu cattle, *Bos indicus* (Canavez et al. 2012). Whereas a total of eight chromosome- and scaffold-level genome assemblies are publicly available for domestic buffaloes, there is currently no genome data available for wild species of the genus *Bubalus*. To fill this gap, a biopsy of a living lowland anoa was used for next-generation sequencing, and a draft genome was assembled *de novo* for comparison to other buffalo genome assemblies available in international databases such as NCBI (National Center for Biotechnology Information) and BIG_GWH (Beijing Institute of Genomics Genome Warehouse database).

## 2. Material & Methods

### 2.1 DNA extraction, library preparation and genome sequencing

A living male adult of lowland anoa, named Yannick, was sampled at the *Ménagerie du Jardin des Plantes* of the Muséum national d’Histoire naturelle (MNHN, Paris, France) (Figure 1). A skin biopsy was performed in 2006 by a veterinary surgeon following protocols approved by the MNHN and in line with ethical guidelines. The same biopsy was previously used to determine its karyotype (2n = 48; FN = 58; Nguyen et al., 2008). DNA was extracted using the DNeasy Blood and Tissue Kit (Qiagen, Hilden, Germany) following the manufacturer’s protocol. DNA quantification was performed with a Qubit^®^ 2.0 Fluorometer with Qubit dsDNA HS Assay Kit (Thermo Fischer Scientific, Walthan, MA, USA). Library preparation and sequencing were conducted at the *Institut du Cerveau et de la Moelle épinière*. The sample was sequenced on a NextSeq^®^ 500 Illumina system generating 2 × 151 bp reads using the NextSeq 500 High Output Kit v2 with 300 cycles and aiming for an insert size of 350 bp.

**Figure 1:**
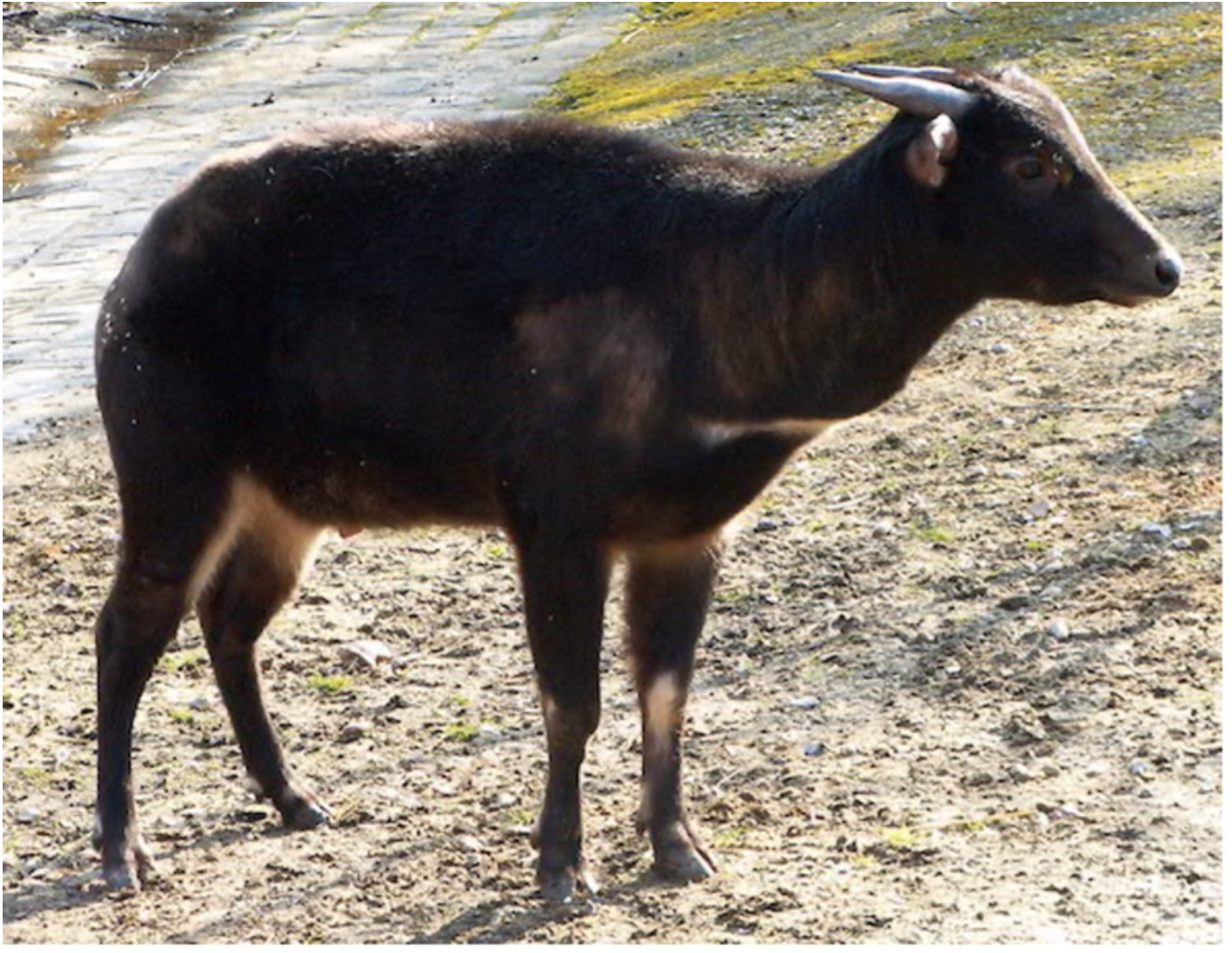
Lowland anoa (*Bubalus depressicornis*) housed at the Mènagerie du Jardin des Plantes (© Alexandre Hassanin - MNHN).

### 2.2 De novo assembly

Data quality was assessed with FastQC v.0.11.5 (https://www.bioinformatics.babraham.ac.uk/projects/fastqc/) and results were collated with MultiQC v1.12 (Ewels et al., 2016). Raw reads were quality trimmed and adapter sequences and contaminants removed with Trimmomatic v.0.36 (Bolger et al., 2014) with the following parameters: “ILLUMINACLIP: TruSeq3 -PE.fa:2:30:10 LEADING:33 TRAILING:3 SLIDINGWINDOW:4:15 MINLEN:36”. Data quality of quality-trimmed reads was re-assessed with FastQC. A *de novo* assembly was performed with MaSuRCA v.3.3.1 (Zimin et al., 2013; Zimin et al., 2017) using recommended parameters for mammalian genomes and paired-end Illumina-only data, as indicated in Zimin et al. (2017). Mean and standard deviation for the Insert size were estimated with an “estimate-insert-size” script (https://gist.github.com/rchikhi/7281991). Paired-end reads were error corrected using QuorUM (Marçais et al., 2015) and assembled into super-reads using a k-mer size of 99, as selected by the MaSuRCA assembler. The super-reads were then assembled into contigs using the CABOG assembler, part of the MaSuRCA pipeline (Zimin et al., 2017), followed by gap closing with the paired-end information (Zimin et al., 2013).

### 2.3 Assembly quality assessment

Genome assemblies publicly available for *Bubalus* and *Syncerus* genera were retrieved from NCBI and BIG_GWH for quality comparison and assessment. The dataset included two assemblies at the chromosome level for the river buffalo (*Bubalus bubalis*) with a coverage of 100x and 572x, four scaffold-level draft assemblies of river buffalo with coverage ranging between 69x and 119x, one chromosome-level assembly of swamp buffalo (*Bubalus kerabau*)with a mean coverage of 65x, and one scaffold-level draft assembly of the African buffalo (*Syncerus caffer*) with 162x coverage. The eight retrieved assemblies were sequenced and assembled with different methods, summarised in Table 1.

**Table 1:**
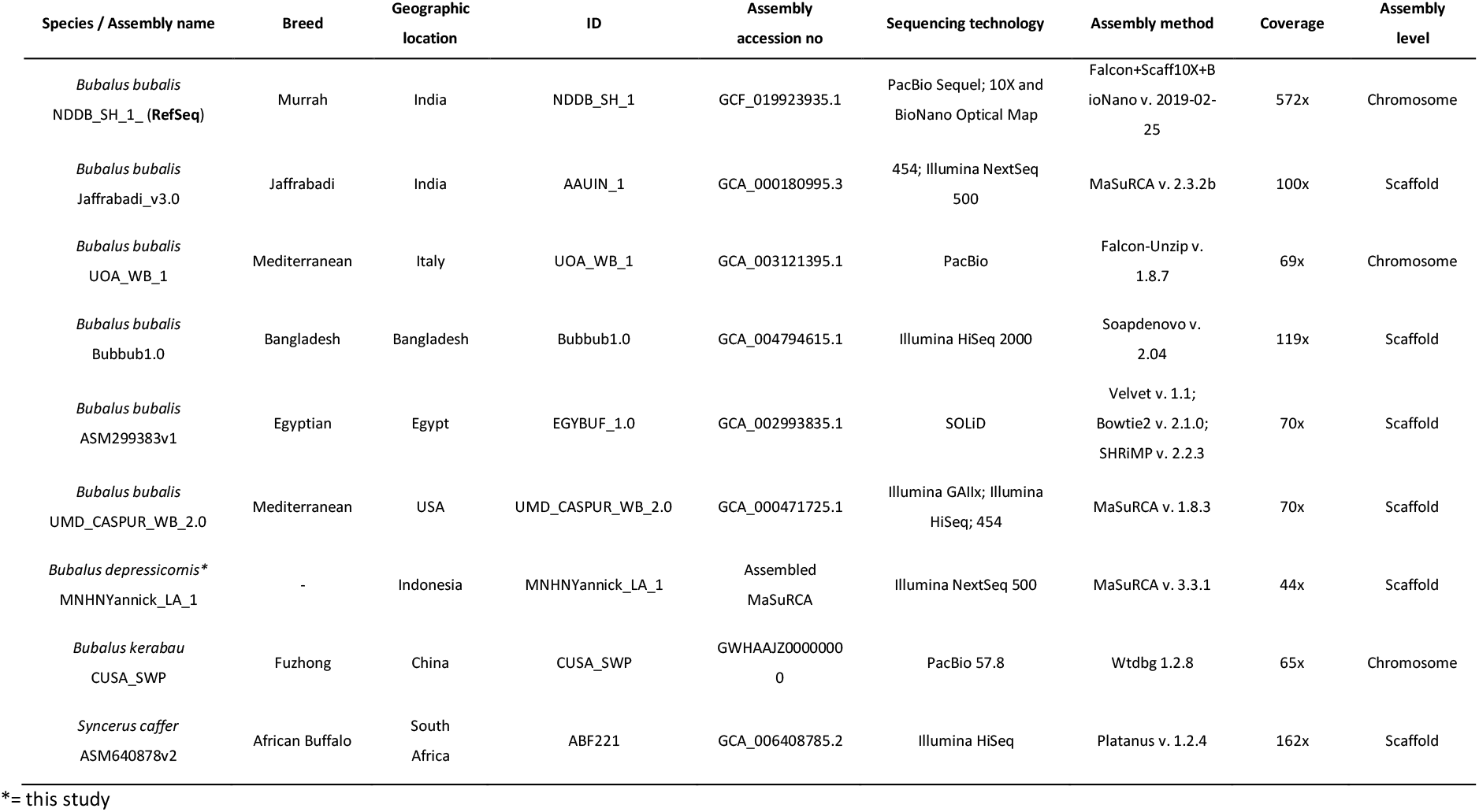
Information regarding genome assemblies available for buffalo species.

The quality of the lowland anoa genome assembly was assessed with QUAST-LG v.5.0.1 (Mikheenko et al., 2018) using the river buffalo NDDB_SH_1 genome assembly (Deng et al., 2016) as a reference. The default parameters for mammalian genomes were used to compare all assemblies in QUAST-LG: “MODE: large, threads: 50, eukaryotic: true, minimum contig length: 3,000, minimum alignment length: 500, ambiguity: one, threshold for extensive misassembly size: 7,000”. All analysed assemblies were aligned to the river buffalo NDDB_SH_1 assembly and results were plotted with Circos v. 0.69.8 (Krzywinski et al., 2009) and Jupiter consistency plots (Chu, 2018).

We adopted two different strategies to evaluate genome completeness. Firstly, genomic features were predicted with the homology-based method by aligning the lowland anoa genome to that of the annotated river buffalo reference genome (NDDB_SH_1 and relative annotations retrieved from NCBI). Secondly, we used a *de novo* gene prediction method with GlimmerHMM v3.0.4 (Majoros et al. 2004). Thirdly, we employed benchmarking against universal single-copy orthologs (BUSCO v5.2.2; Manni et al. 2021) using the mammalia_odb10 dataset (19/02/2021, number of genomes: 24, number of BUSCOs: 9226) from OrthoDB (Kriventseva et al. 2019) and compared to other buffalo genome assemblies already deposited on NCBI and BIG_GWH (Table 1).

### 2.4 Repeats and gene annotation

Repetitive regions in the lowland anoa genome were identified, annotated and masked with RepeatMasker v.4.1.2-p1 (Tarailo-Graovac and Chen, 2009). Firstly, a *de novo* repeat library was constructed from the genome assembly with RepeatModeler v.2.0.2a. RepeatMasker was used with default parameters to produce a homolog-based repeat library and mask the genome’s repetitive regions. The scripts *“calcDivergenceFromAlign.pl”* and *“createRepeatLandscape.pl”* were used to calculate the Kimura divergence values and to plot the resulting repeat landscape. The repeat landscape of *Bos taurus* was retrieved from the RepeatMasker database for visual comparison.

## 3. Results & Discussion

### 3.1 Whole-genome sequencing and data QC

Whole-genome sequencing generated 991,437,058 paired-end reads with a length of 151 bp. Quality trimming removed 46,616,722 low quality, adapter-contaminated and PCR-duplicated reads, representing approximately 0.5% of the total reads. A total of 944,820,336 clean paired-end reads were generated, covering the lowland anoa genome with an estimated 56x depth based on a genome size of 2.56 Gbp. Estimation of insert size using in-house script returned a mean of 377 and a standard deviation of 83.

### 3.2 De novo assembly quality metrics

The final lowland anoa genome assembly generated here contained 103,135 contigs, the largest being 337.39 kbp long, an N50 of 38.73 kbp and an L50 of 19.83 kbp (Table 2). Total length was 2.56 Gbp with a mean coverage of 44x, and GC content was 41.74%, in agreement with other published assemblies (between 41.60% and 41.92%, Table 3). When aligned to the NDDB_SH_1 genome assembly, the fraction of the anoa genome assembly was 95.41%, a value comparable to other buffalo genome assemblies (Figure 2), with a total alignment length of 2,515,453,843 bp. A total of 886 contigs could not be aligned to the river buffalo genome assembly, whilst 8,085 contigs were only partially aligned, resulting in a total unaligned length of 45,224,171 bp, which reflects the discrepancy between the total length of the lowland anoa genome and the total aligned length to the reference river buffalo genome assembly. Partially aligned and unaligned contigs could have resulted from structural variations between the lowland anoa and the reference river buffalo assembly, such as large INDELS (insertion/deletions), as well as repetitive regions and/or alternative haplotypes causing assembly errors. The nature of short-read technology causes difficulties in characterising genomic regions such as telomeres, centromeres, repetitive and highly heterochromatic regions (Johnson et al. 2005; Low et al. 2019; Weissensteiner and Suh 2019), which are notoriously difficult to assemble and could be better resolved with long-read sequencing.

**Figure 2:**
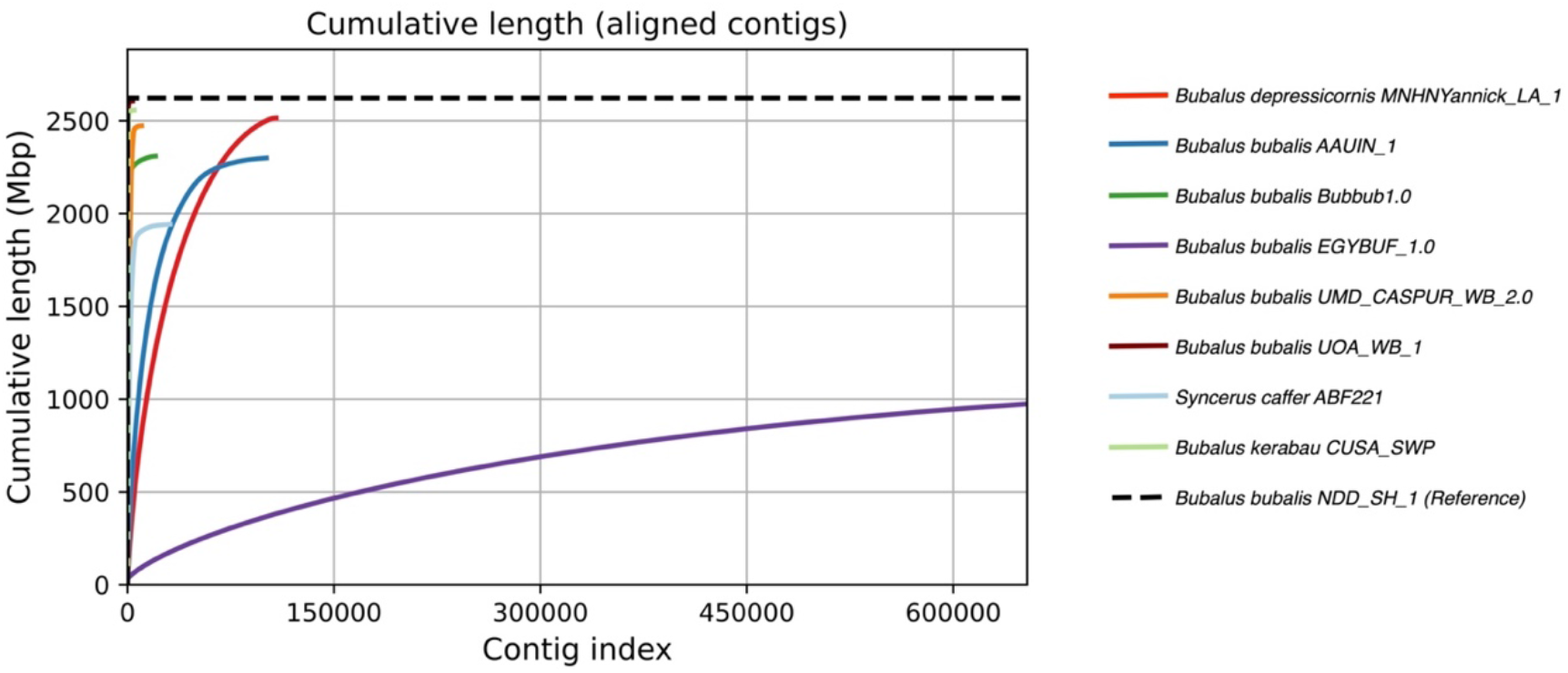
Cumulative length of aligned contigs of the lowland anoa (red line) against the river buffalo NDDB_SH_1 reference genome assembly (dashed line) and compared to other buffalo genome assemblies available on NCBI.

**Table 2:**
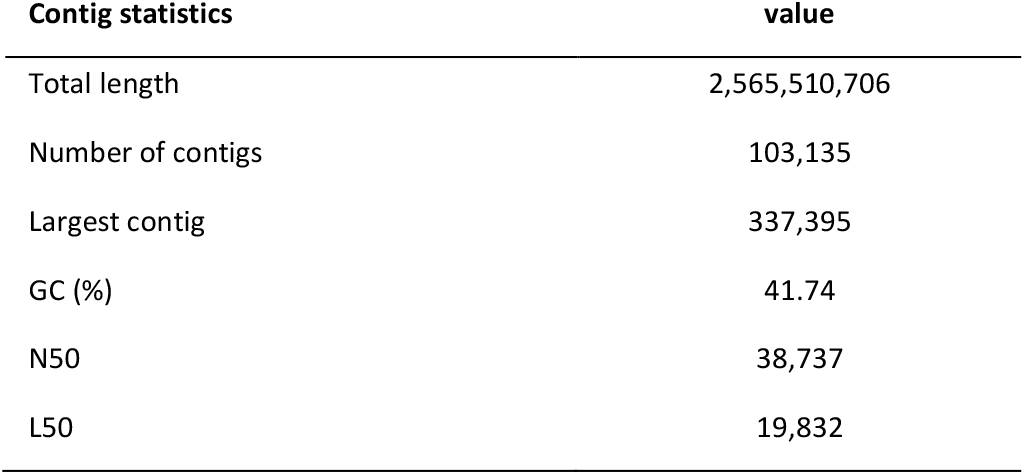
Draft assembly statistics of the lowland anoa genome

**Table 3:**
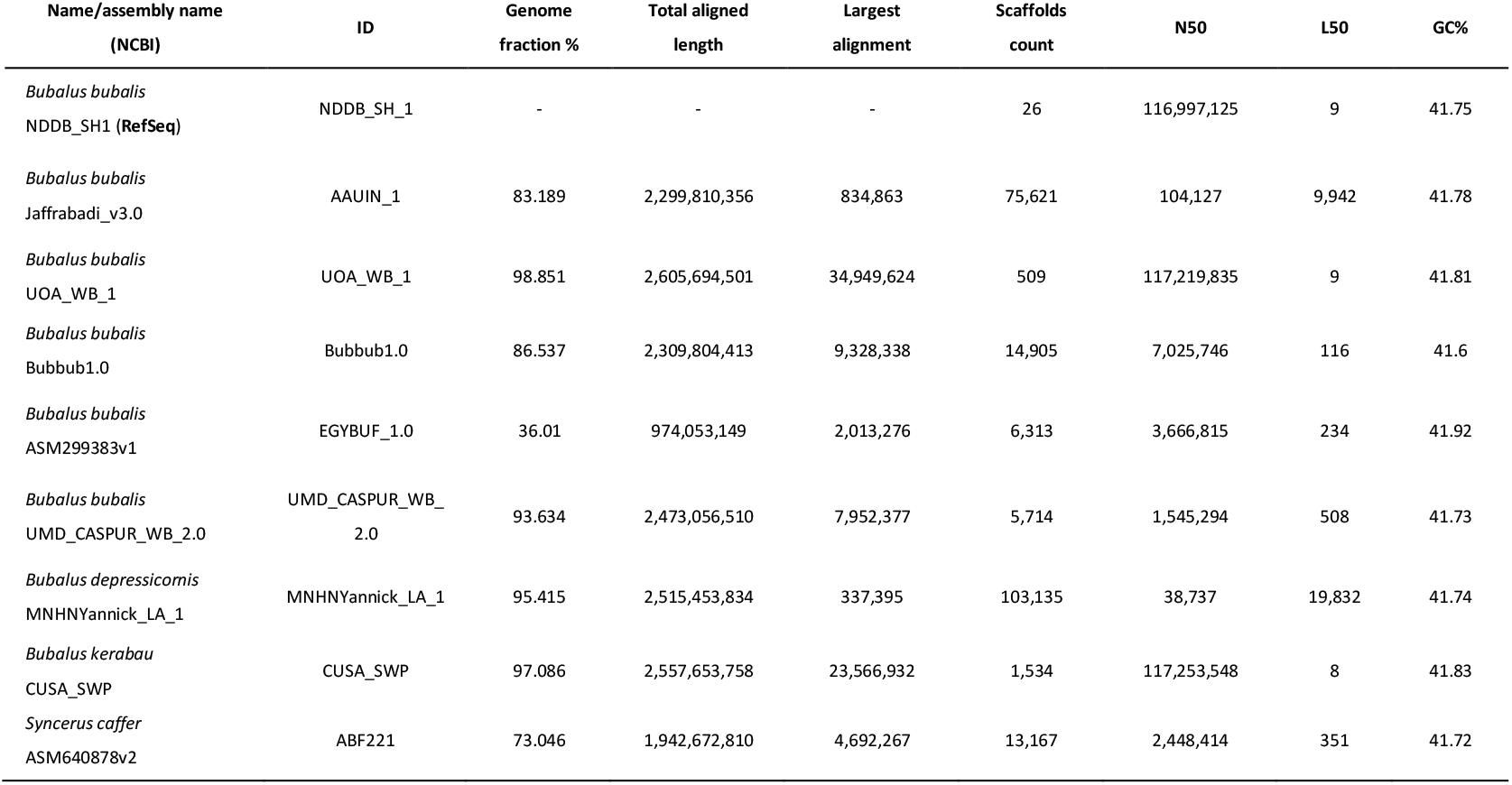
Comparison of assembly quality metrics of the lowland anoa (*Bubalus depressicornis*) and other buffalo assemblies.

The lowland anoa genome assembly has a modest N50 compared to other buffalo genome assemblies (Table 3), indicating lower levels of contiguity, which is expected due to the short-read output of Illumina sequencing technology (read length = 151 bp). Additionally, repeat analysis revealed that 42.12% of the lowland anoa genome is composed of repetitive regions. This, coupled with low sequence coverage, sequencing and assembly errors, causes breaks in the assembly contiguity (Gnerre et al., 2011; Low et al., 2019). This is apparent even in high-quality chromosome-level genome assemblies that use multiple sequencing libraries and multiple sequencing technologies, such as the previous human genome assembly GRCh38, which contained hundreds of gaps (International Human Genome Sequencing Consortium 2004). In addition, the chromosome-level genome assemblies retrieved from NCBI (NDDB_SH_1, UOA_WB_1) were sequenced using multiple insert size libraries and sequencing technologies and were intensively verified with multiple methods such as optical mapping, Hi-C and RH (Deng et al., 2016; Low et al., 2019).

Moreover, quality metrics of publicly available assemblies are usually limited to reporting N50 and L50 values, which represent the shortest contig length needed to cover 50% of the total assembly size, and the number of contigs whose cumulative length covers 50% of the total assembly size, respectively (Bradnam et al., 2013). Such metrics are often used to compare and evaluate performances of the ever-growing assembly and annotation methods and software (Manchanda et al., 2020). However, we hereby show that reporting N50 and L50 metrics exclusively can be misleading, as they only provide a standard measure of assembly contiguity whilst omitting information such as gene content and completeness, as well as assembly correctness. Furthermore, N50 values can be artificially raised by deliberately excluding short contigs from analyses and by the presence of undetermined nucleotides (Ns) linking the scaffolded contigs (Gurevich et al. 2013). Therefore, to assess the quality of the lowland anoa genome assembly, we generated conventional N50 and L50 metrics and also determined genome completeness in terms of gene content and genome correctness by comparing our assembly to a chromosome level genome assembly of the river buffalo (*Bubalus bubalis*). Additionally, a swamp buffalo (*Bubalus kerabau*, CUSA_SWP) and a more distantly related African buffalo species (*Syncerus caffer*, ABF221) were also included in our comparison.

Regardless of the modest N50 value, the lowland anoa genome assembly is in good agreement with the NDDB_SH_1 assembly, with 95.91% of contigs correctly mapped to the 25 reference chromosomes of the river buffalo and fewer misassembled blocks compared to other draft assemblies (Figure 3). The genome assembly of the Egyptian river buffalo (EGYBUF_1.0) had an abnormally high number of misassembled blocks with respect to the reference genome, followed by the genome assembly of a female Italian river buffalo (UOA_WB_1). To investigate this, misassemblies and structural variation metrics were computed in QUAST-LG (Table 4). The Egyptian river buffalo assembly (EGYBUF_1.0) showed the highest number of mismatches and the highest number of Ns, followed by the Jaffrabadi river buffalo (AAUIN_1). The genome assembly of the African buffalo (*S. caffer*, ABF221) showed a larger number of mismatches (Table 4), but this can be explained by the higher sequence divergence between *Syncerus* and *Bubalus*, as the two genera have separated in the Late Miocene (Hassanin et al., 2012). Misassemblies and structural variation metrics could not explain the misassembled blocks of the UOA_WB_1 assembly observed in the Circos plot of Figure 3. However, some of these misassembled blocks could be due to unplaced contigs. To investigate this, the UOA_WB_1 assembly was aligned to the NDDB_SH_1 reference to generate Jupiter consistency plots. When using the largest 26 contigs of the UOA_WB_1 assembly to cover 100% of the reference river buffalo genome, an almost perfect level of synteny was observed (Figure 4a). Although this result was expected for genomes of the same species, it also indicates a good level of assembly quality in terms of correctness. However, when including all 509 contigs of the UOA_WB_1 assembly, several misassembled regions were observed (Figure 4b). Three non-exclusive hypotheses can be advanced to interpret this result: possible genomic rearrangements, genome assembly errors, and repetitive regions. Whether the results of the consistency plots are due to the factors mentioned above or other factors, such as contamination, remains speculative. Nevertheless, the results of the quality metric comparison conducted here further indicate the unreliability of using exclusively N50 and L50 metrics when assessing assembly quality. Instead, contiguity metrics should be supplemented with genome completeness and correctness metrics.

**Figure 3:**
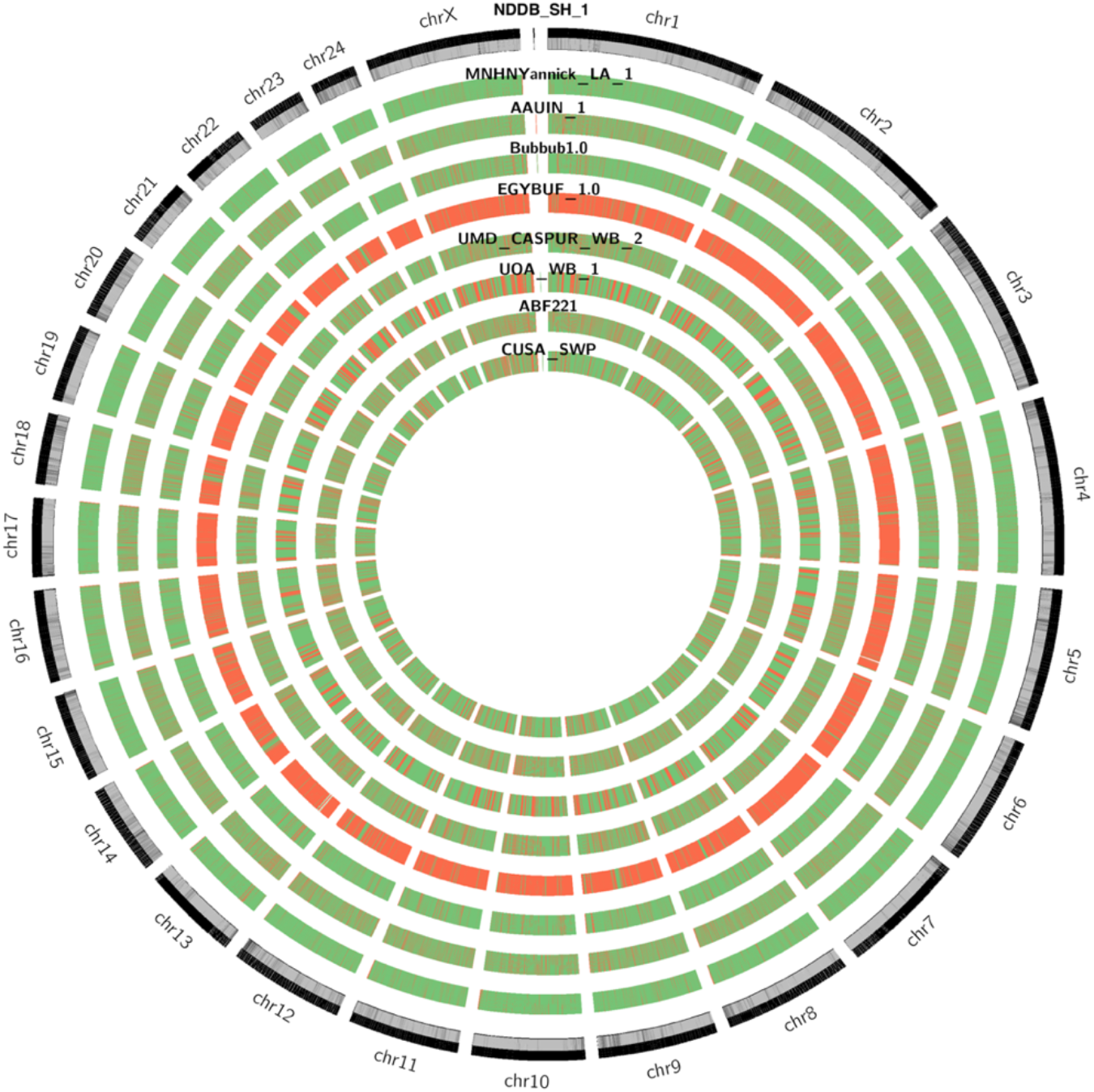
Circos plot of scaffolds mapped to NDD_SH_1 reference genome assembly (*Bubalus bubalis)*. Outer circle represents reference sequence with GC% heatmap (0%= white, 69%= black). Inner circles represent assembly tracks, with heatmap representing correct contigs (green) and misassembled blocks (red).

**Figure 4:**
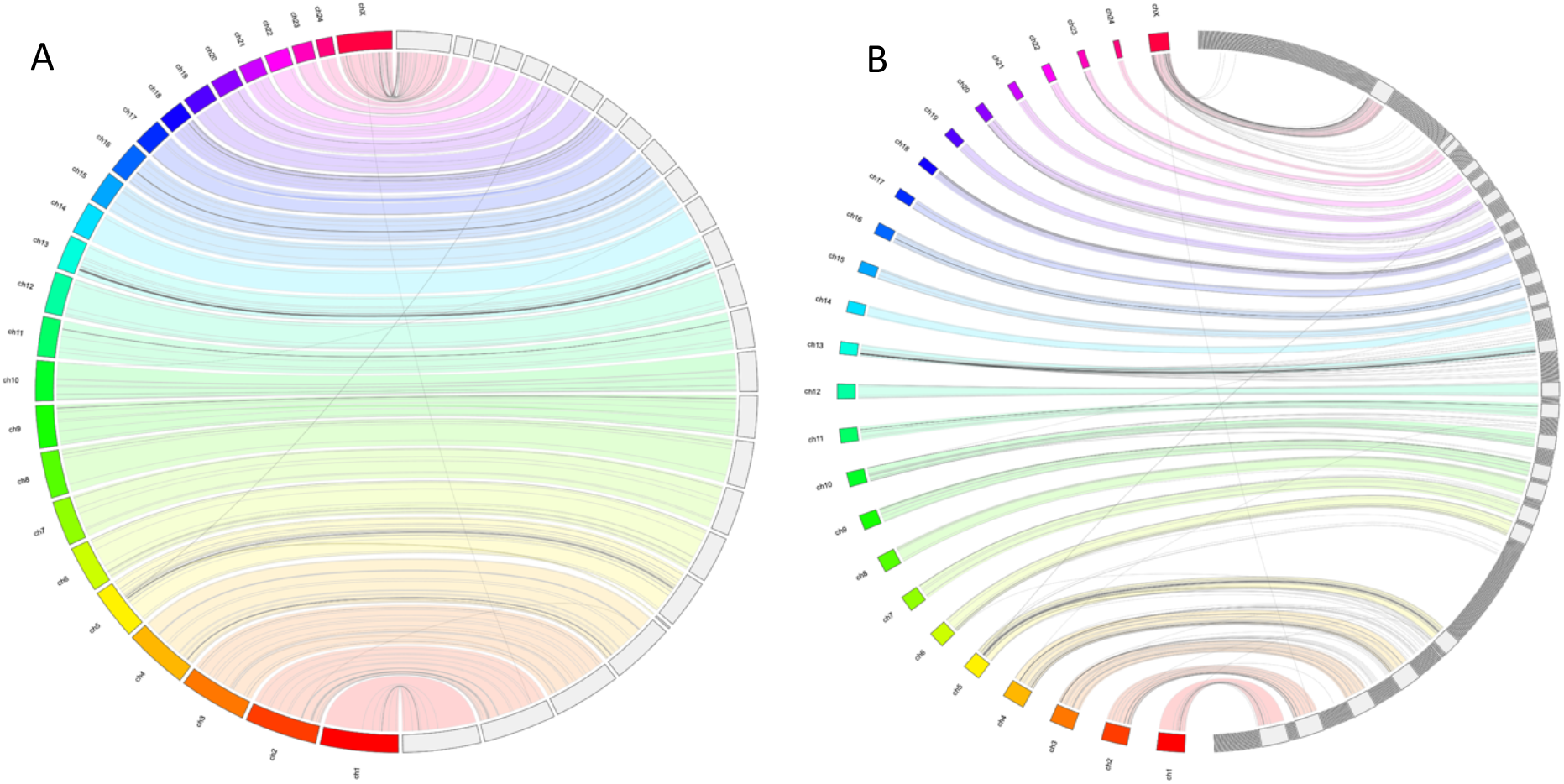
Jupiter consistency plot showing alignment between the river buffalo genome assemblies UO_AWB_1 and NDDB_SH_1. The left of the plots shows the numbered NDDB_SH_1 chromosomes. The right of the plots shows (A) the 26 longest contigs of the UOA_WB_1 assembly needed to cover 100% of the reference genome, and (B) all the 509 contigs of the UO_AWB_1 assembly. Coloured bands represent synteny between the genomes. Lines represent genomic rearrangements, break points in the scaffolds or assembly errors. Absence of lines connecting the UO_AWB_1 blocks to the NDDB_SH_1 chromosomes indicates contigs that could not be aligned to the reference.

**Table 4:** QUAST-LG statistics of all buffalo assemblies with respect to the river buffalo NDDB_SH_1 reference.

|  | *B. depressicornis*<br>MNHNYannick_LA_1 | *B. bubalis*<br>AAUIN_1 | *B. bubalis*<br>Bubbub1.0 | *B. bubalis*<br>EGYBUF_1.0 | *B. bubalis*<br>UMD_CASPUR_WB_2.0 | *B. bubalis*<br>UOA_WB_1 | *B. kerabau*<br>CUSA_SWP | *S. caffer*<br>ABF221 |
| --- | --- | --- | --- | --- | --- | --- | --- | --- |
| **Misassemblies** | 4,949 | 19,238 | 3,561 | 131 | 4,040 | 1,724 | 2,111 | 6,565 |
| **Relocations** | 1,447 | 13,540 | 2,761 | 85 | 1,434 | 1,051 | 1,199 | 3,397 |
| **Translocations** | 3,203 | 4,714 | 757 | 10 | 2,569 | 647 | 896 | 3,032 |
| **Inversions** | 299 | 984 | 43 | 36 | 37 | 26 | 16 | 136 |
| **Misassembled contigs** | 4,550 | 15,988 | 1,049 | 45 | 1,943 | 255 | 533 | 1,727 |
| **Misassembled contigs length** | 159,179,266 | 1,334,096,556 | 2,506,642,146 | 55,459,162 | 1,891,377,139 | 2,639,940,877 | 2,594,120,526 | 2,486,555,687 |
| **Local misassemblies** | 7,014 | 73,267 | 241,261 | 6,933 | 7,100 | 4,870 | 9,940 | 435,454 |
| **Possible TEs** | 164 | 874 | 886 | 10 | 544 | 136 | 158 | 654 |
| **Unaligned mis. contigs** | 287 | 2,378 | 548 | 2,522 | 63 | 104 | 381 | 1,324 |
| **Unaligned contigs** | 886 + 8,085<br>partial | 2,555 + 57,865<br>partial | 297 + 7,280<br>partial | 2,806 + 3,472<br>partial | 182 + 3,290<br>partial | 1 + 416<br>partial | 140 + 1110<br>partial | 900 + 7,314<br>partial |
| **Unaligned length** | 45,224,171 | 596,227,806 | 299,544,303 | 1,673,093,194 | 82,826,374 | 49,291,638 | 51,316,520 | 779,611,955 |
| **Genome fraction (%)** | 95.415 | 83.189 | 86.537 | 36.01 | 93.634 | 98.851 | 97.086 | 73.046 |
| **Duplication ratio** | 1.007 | 1.425 | 1.076 | 1.36 | 1.034 | 1.005 | 1.013 | 1.045 |
| **Mismatches** | 16,233,421 | 19,654,061 | 23,375,163 | 17,890,296 | 10,863,130 | 10,118,782 | 15,844,866 | 114,608,168 |
| **Indels** | 1,578,224 | 746,243 | 705,955 | 6,440,610 | 1,136,878 | 1,400,310 | 1,534,735 | 2,128,964 |
| **Indels length** | 12,654,316 | 56,163,406 | 24,209,936 | 35,356,432 | 24,745,254 | 23,411,739 | 33,123,824 | 18,236,722 |
| **Mismatches per 100 kbp** | 649 | 901 | 1,030 | 1,895 | 442 | 390 | 622 | 5,983 |
| **Indels per 100 kbp** | 63 | 34 | 31 | 682 | 46 | 54 | 60 | 111 |
| **indels (<= 5 bp)** | 1,297,998 | 598,354 | 515,830 | 5,758,980 | 893,802 | 1,227,309 | 1,269,689 | 1,641,754 |
| **indels (> 5 bp)** | 280,226 | 147,889 | 190,125 | 681,630 | 243,076 | 173,001 | 265,046 | 487,210 |
| **N's** | 493,027 | 850,098,824 | 138,209,713 | 328,128,682 | 73,946,361 | 373,500 | 22,116,406 | 59,283,755 |
| **N's per 100 kbp** | 19.22 | 22,942 | 5,040.03 | 11,097 | 2,820.18 | 14.06 | 840.50 | 2,131.26 |

### 3.3 Genomic features, gene prediction and annotation

Homology and *de novo* gene predictions performed on the lowland anoa genome assembly were in agreement with each other and indicated a good level of genome completeness. Results were comparable to other published genome assemblies (Tables 5 and 6), and an improvement over the Bangladeshi river buffalo (Bubbub_1.0), the Egyptian river buffalo (EGYBUF_1.0) and Mediterranean river buffalo (UMD_CASPUR_WB_2.0) assemblies.

Interestingly, these three assemblies showed higher contiguity (N50) than the draft assembly of the lowland anoa, further indicating the unreliability of using exclusively N50 and L50 metrics when assessing genome assembly quality.

Out of the 1,921,249 genomic features annotations of the reference assembly NDDB_SH_1, homology prediction identified 1,815,794 (94.51%) complete and 69,929 (3.63%) partial features in the lowland anoa genome assembly, which is comparable to other published assemblies (Figure 5), indicating a good level of genome completeness. GlimmerHMM *de novo* predicted 1,027,469 unique genomic features (mRNA and coding sequences, CDS), which is an improvement over some of the water buffalo assemblies used for quality comparison (Table 5). Homology-based gene prediction identified 32,393 genes in the lowland anoa genome assembly, representing 97.14% of the genes annotated in NDDB_SH_1 (n= 33,348). Of these, 59.11% (19,148) were complete and 40.88% (13,245) were partial, probably reflecting the level of fragmentation of the lowland anoa genome assembly. Nevertheless, the total number of genes predicted still represents an improvement over some of the compared assemblies (Table 6).

**Figure 5:**
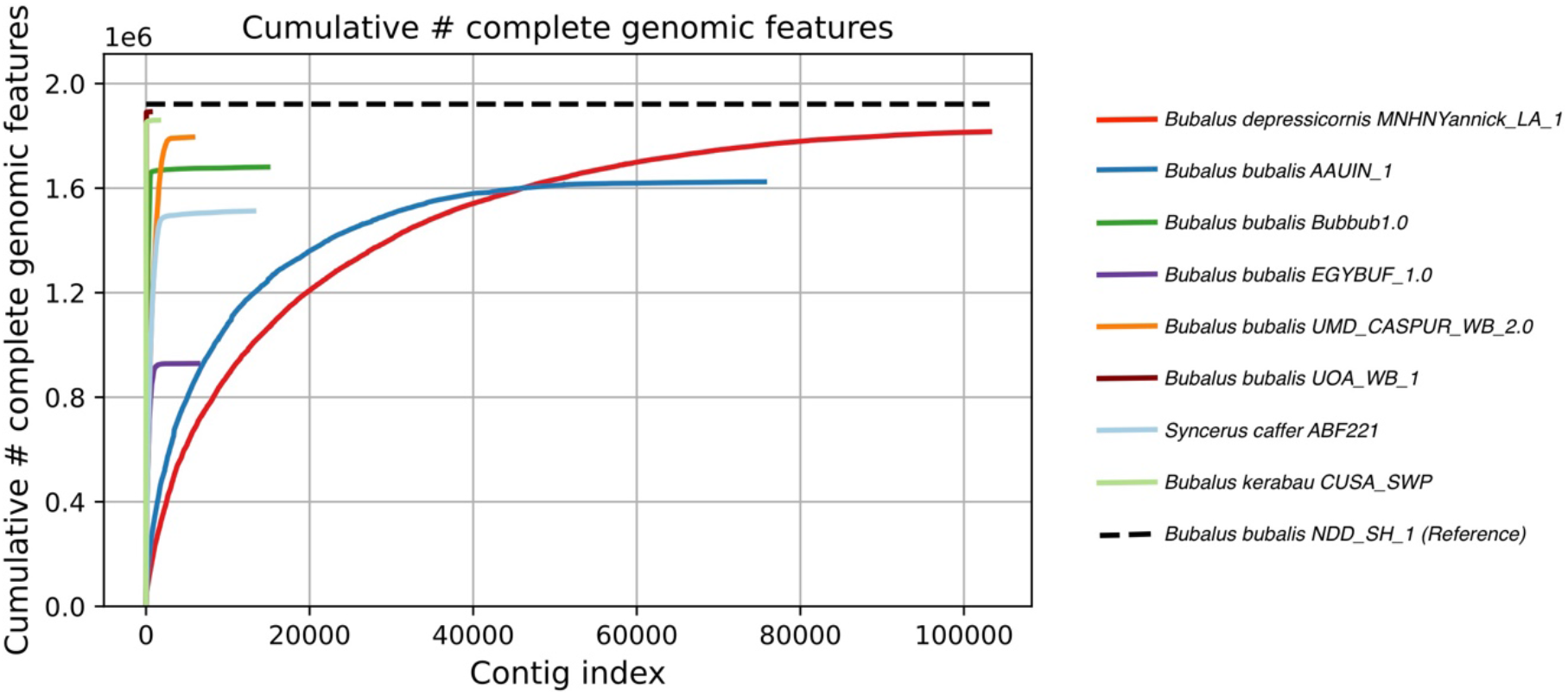
Complete genomic features identified in the lowland anoa assembly and compared to other assemblies using the river buffalo (*Bubalus bubalis*) NDD_SH1 reference sequence and annotations.

**Table 5:** Gene features (CDS and mRNA) predicted with GlimmerHMM

| Name/assembly name<br>(NCBI) | ID | predicted<br>gene<br>features<br>(unique) | predicted gene<br>features (>= 0 bp) | predicted gene<br>features (>= 300 bp) | predicted gene<br>features (>= 1500<br>bp) | predicted gene<br>features (>= 3,000 bp) |
| --- | --- | --- | --- | --- | --- | --- |
| *Bubalus bubalis*<br>Jaffrabadi_v3.0 | AAUIN_1 | 1,065,654 | 1,087,174 + 1,214 part | 719,235 + 911 part | 129,801 + 19 part | 24,579 + 7 part |
| *Bubalus bubalis*<br>UOA_WB_1 | UOA_WB_1 | 1,055,791 | 1,059,972 + 21 part | 762,464 + 17 part | 154,594 + 0 part | 29,659 + 0 part |
| *Bubalus bubalis*<br>Bubbub1.0 | Bubbub1.0 | 948,732 | 958,663 + 101 part | 655,839 + 73 part | 136,045 + 4 part | 27,867 + 1 part |
| *Bubalus bubalis*<br>ASM299383v1 | EGYBUF_1.0 | 826,048 | 826,155 + 69 part | 530,835 + 37 part | 96,365 + 0 part | 16,243 + 0 part |
| *Bubalus bubalis*<br>UMD_CASPUR_WB_2.0 | UMD_CASPUR_WB_2.0 | 963,177 | 964,473 + 138 part | 669,508 + 117 part | 134,780 + 5 part | 26,448 + 2 part |
| *Bubalus depressicornis*<br>MNHNYannick_LA_1 | MNHNYannick_LA_1 | 1,027,469 | 1,023,163 + 5,278 part | 702,282 + 4,582 part | 131,966 + 204 part | 24,994 + 37 part |
| *Bubalus kerabau*<br>CUSA_SWP | CUSA_SWP | 1,042,862 | 1,046,662 + 87 part | 752,170 + 70 part | 151,809 + 10 part | 29,488 + 6 part |
| *Syncerus caffer*<br>ASM640878v2 | ABF221 | 1,061,091 | 1,064,542 + 229 part | 750,719 + 171 part | 150,033 + 10 part | 29,460 + 1 part |

**Table 6:** Genes predicted with homology-based prediction method.

| Name/assembly name<br>(NCBI) | ID | Genes | Partial genes | Total | % Of reference's<br>annotated genes<br>(n= 33,348) |
| --- | --- | --- | --- | --- | --- |
| *Bubalus bubalis*<br>Jaffrabadi_v3.0 | AAUIN_1 | 10,804 | 20,895 | 31,699 | 95.05 |
| *Bubalus bubalis*<br>UOA_WB_1 | UOA_WB_1 | 30,810 | 1,955 | 32,765 | 98.25 |
| *Bubalus bubalis*<br>Bubbub1.0 | Bubbub1.0 | 11,039 | 20,983 | 32,022 | 96.02 |
| *Bubalus bubalis*<br>ASM299383v1 | EGYBUF_1.0 | 1,345 | 23,770 | 25,115 | 75.31 |
| *Bubalus bubalis*<br>UMD_CASPUR_WB_2.0 | UMD_CASPUR_WB_2.0 | 18,656 | 13,271 | 31,927 | 95.74 |
| *Bubalus depressicornis*<br>MNHNYannick_LA_1 | MNHNYannick_LA_1 | 19,148 | 13,245 | 32,393 | 97.14 |
| *Bubalus kerabau*<br>CUSA_SWP | CUSA_SWP | 28,349 | 3,419 | 31,768 | 95.26 |
| *Syncerus caffer*<br>ASM640878v2 | ABF221 | 8,763 | 21,575 | 30,338 | 90.97 |

When predicting mammalian orthologs with BUSCO, the lowland anoa genome assembly contained 6,556 (71.1%) complete BUSCOs, of which 6,412 (69.5%) were single-copy and 144 (1.6%) were duplicated. The number of fragmented BUSCOs was 1,076 (11.7%), whilst 1,594 (17.2%) were missing. The BUSCO results indicate an acceptable level of genome completeness (<70%, Simão et al., 2015) for downstream analyses for the anoa genome assembly, and a slight improvement over the Egyptian river buffalo assembly (EGYBUF_1.0, Figure 6).

**Figure 6:**
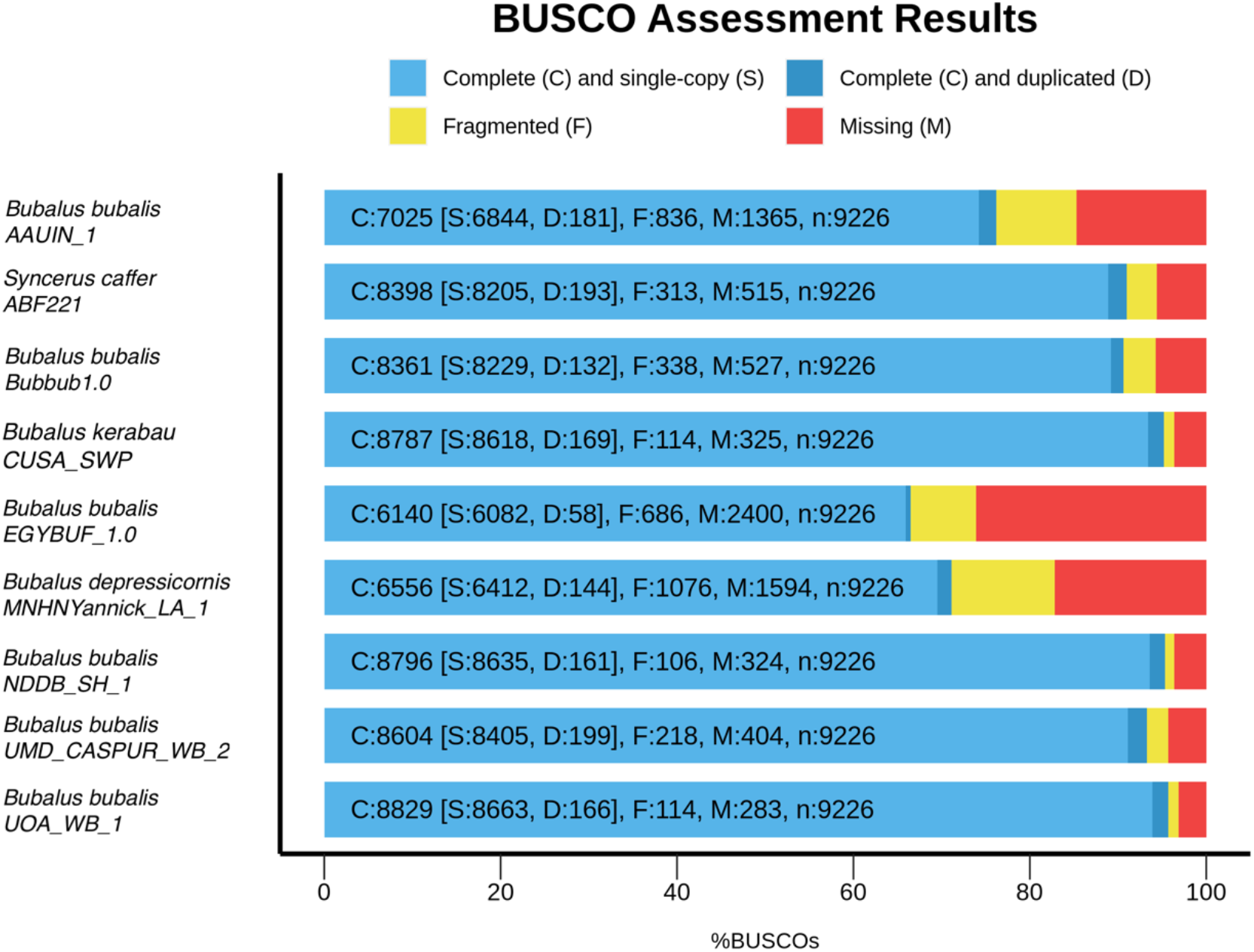
BUSCO results of the genome assembly of the lowland anoa (*Bubalus depressicornis*) compared to other publicly available buffalo genome assemblies.

Mammalian genomes contain large families of repeats (Goodier and Kazazian, 2008), such as long interspersed nuclear elements (LINEs), short interspersed nuclear elements (SINEs), and long-terminal repeats (LTRs). RepeatMasker revealed that 42.12% of the lowland anoa genome is composed of repetitive regions (Table 7), which is comparable to data previously published for genome assemblies of river buffalo and other bovids (Deng et al., 2016; Low et al., 2019; Mintoo et al,. 2019; El-Khishin et al., 2020). Results also agree with the repetitive content in the cattle genome (Figure 7b). Both lowland anoa and cattle genomes showed two waves of repeat expansion in their repeat landscape (Figure 7a and 7b), suggesting a shared inheritance of such repeats. In the lowland anoa, the LINEs were more abundant, representing 30.04% of the repeats, followed by LTRs representing 3.10% and SINEs representing 1.03% (Table 7).

**Figure 7:**
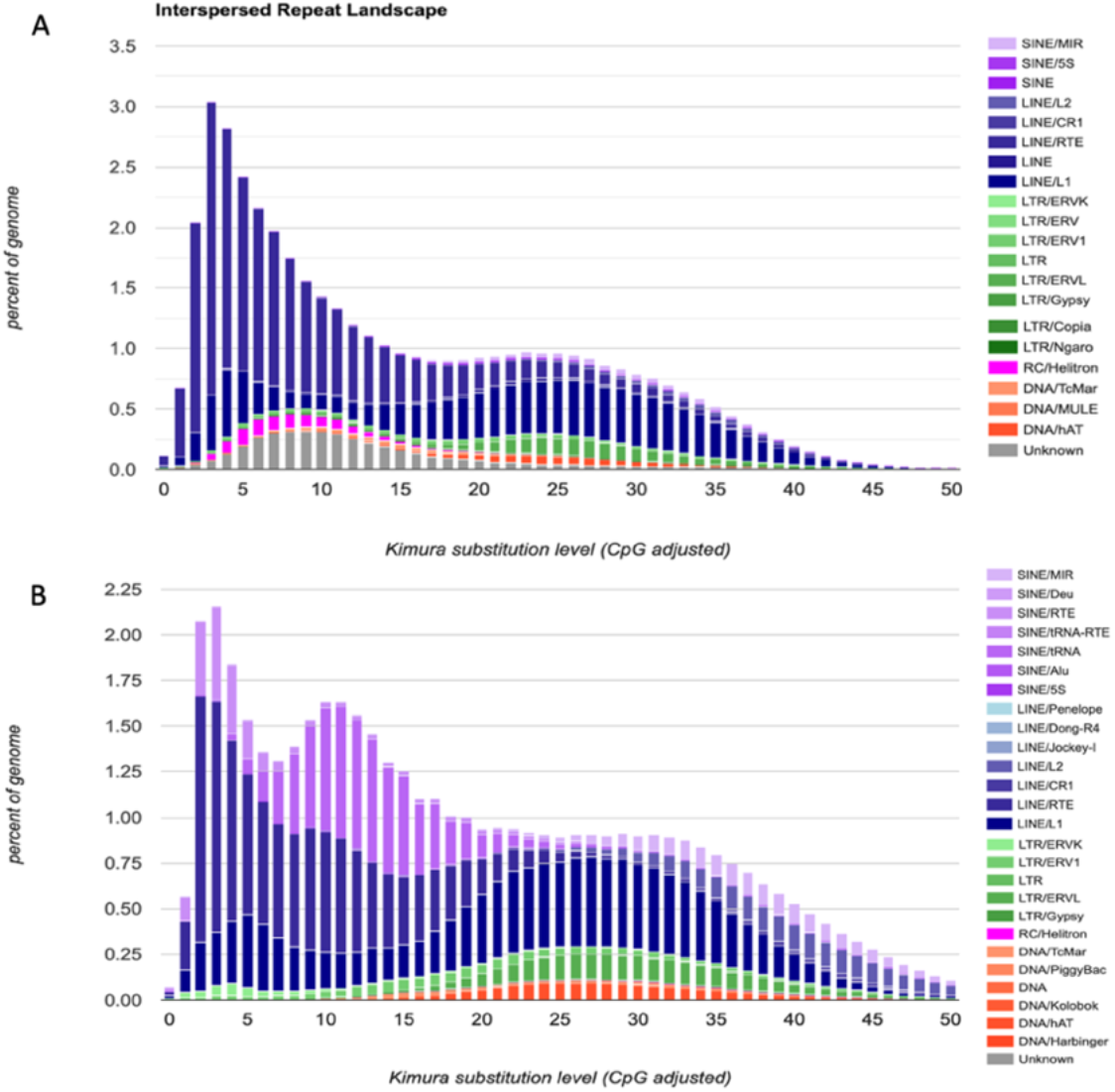
Interspersed Repeat Landscape of (**A**) the lowland anoa genome assembled in this study and (**B**) *Bos taurus*.

**Table 7:** Repeat sequence composition of the lowland anoa genome.

| Family | Copy number of elements | Length occupied (bp) | % Genome |
| --- | --- | --- | --- |
| SINEs | 296,064 | 26,945,915 | 1.03% |
| LINEs | 2,864,468 | 786,815,034 | 30.04% |
| LINE1 | 1,203,360 | 282,366,346 | 10.78% |
| LINE2 | 101,415 | 13,911,301 | 0.53% |
| RTE/Bov-B | 1,461,651 | 481,114,012 | 18.37% |
| LTR elements | 362,123 | 81,208,077 | 3.10% |
| DNA transposon | 255,003 | 38,433,935 | 1.47% |
| Small RNA | 139,586 | 14,174,190 | 0.54% |
| Satellites | 269 | 52,169 | 0.00% |
| Simple repeats | 500,363 | 20,187,327 | 0.77% |
| Low complexity | 81,685 | 3,956,146 | 0.15% |
| Unclassified | 611,789 | 100,086,577 | 3.82% |
| Total |  |  | 42.12% |

## 4. Conclusion

To date, whole-genome sequencing has allowed identification of variants involved in domestication and genetic improvement for several livestock species (Zimin et al., 2009; Canavez et al., 2012; Li et al., 2020; Rosen et al., 2020). However, the lack of wild buffalo genomes hinders further analyses addressing functional and evolutionary aspects of this group, as well as possible conservation efforts. The draft genome assembly of the lowland anoa reported here is expected to contribute to this gap in data availability, as this is the first draft genome assembly for wild Asian buffaloes. Furthermore, we showed that short-read Illumina sequencing data can still provide a cost-effective way of sequencing mammalian genomes to an adequate level of completeness for downstream comparative analyses.

## Data availability

The genome assembly of the lowland anoa is available on NCBI under accession XXXXXXXXXXXXX. The raw data is available on the Sequence Read Archive (SRA) on NCBI under accession XXXXXXXXXX (under embargo until review).

## Acknowledgements

We thank the people of the *Ménagerie du Jardin des Plantes* who helped to collect the biopsy of the lowland anoa used in this study: Norin Chai, Gerard Dousseau, Christelle Hano, Abderrahmane Latreche, Claire Rejaud, Roland Simon, and Rudy Wedlarski. The authors would like to thank Huw Jones for the proofreading of the manuscript.

